# Crucial role of ppGpp in the resilience of *Escherichia coli* to growth disruption

**DOI:** 10.1101/2020.08.31.276709

**Authors:** Clément Patacq, Nicolas Chaudet, Fabien Letisse

## Abstract

Bacteria grow in constantly changing environments that can suddenly become completely deleted in essential nutrients. The stringent response, a rewiring of the cellular metabolism mediated by the alarmone (p)ppGpp, plays a crucial role in adjusting bacterial growth to the severity of the nutritional insult. The ability of (p)ppGpp to trigger a slowdown of cell growth or induce bacterial dormancy has been widely investigated. However, little is known about the role of (p)ppGpp in promoting growth recovery after severe growth inhibition. In this study, we performed a time-resolved analysis of (p)ppGpp metabolism in *Escherichia coli* as it recovered from a sudden slowdown in growth. Results show that *E. coli* recovers by itself from the growth disruption provoked by the addition of serine hydroxamate, the serine analogue that we used to induce the stringent response. Growth inhibition was accompanied by a severe disturbance of metabolic activity and more surprisingly, by a transient overflow of valine and alanine. Our data also show that ppGpp is crucial for growth recovery since in the absence of ppGpp, *E. coli*’s growth recovery was slower. In contrast, an increased concentration of pppGpp was found to have no significant effect on growth recovery. Interestingly, the observed decrease in intracellular ppGpp levels in the recovery phase correlated with bacterial growth and the main effect involved was identified as growth dilution rather than active degradative process. This report thus significantly expands our knowledge of (p)ppGpp metabolism in *E. coli* physiology.

**IMPORTANCE:** The capacity of microbes to resist and overcome environmental insults, know as resilience, allows them to survive in changing environments but also to resist antibiotic and biocide treatments, immune system responses. Although the role of the stringent response in bacterial resilience to nutritional insults has been well studied, little is known about its importance in the ability of the bacteria to not just resist but also recover from these disturbances. To address this important question, we investigated growth disruption resilience in the model bacterium *Escherichia coli* and its dependency on the stringent response alarmone (p)ppGpp by quantifying ppGpp and pppGpp levels as growth was disrupted and then recovered. Our findings may thus contribute to understanding how ppGpp improves *E. coli*’s resilience to nutritional stress and other environmental insults.

## INTRODUCTION

As single-cell organisms, bacteria face constant changes in their direct physico-chemical and nutritional environments. To overcome these disturbances, bacteria have developed adaptive properties that allow them to survive, grow, and eventually evolve. Depletion of external nutrients is one of the most serious insults for these organisms because they have very little internal storage and the fact that their ability to rapidly modulate metabolic functions is key to their survival. A central component of this metabolic adaptation to nutrient stress is the stringent response, a pleiotropic mechanism in bacteria that coordinates growth and nutrient availability (1) and affects a wide range of cellular processes (2). The stringent response is mediated by the accumulation of guanosine tetra- and pentaphosphate (guanosine 3′,5′-bis(diphosphate) and guanosine 3′-diphosphate, 5′-triphosphate), collectively known as (p)ppGpp, which act as second messengers to fundamentally reprogram cellular physiology from rapid growth in rich nutritional environment to survival and adaptation when nutrients become scarce (3, 4). (p)ppGpp also plays other important roles in the regulation of bacterial virulence (5), survival during host invasion (6), and antibiotic resistance and persistence (7–9). Intracellular levels of (p)ppGpp are controlled by RSH (RelA-SpoT Homolog) enzymes, whose name derives from the (p)ppGpp synthetase RelA and the (p)ppGpp synthetase/hydrolase SpoT in *Escherichia coli*, where (p)ppGpp was originally detected (10).

In this bacterium, it has been established for decades that the reaction to amino acid limitation is a RelA-mediated-stringent response (3). RelA is a GDP/GTP pyrophosphokinase that, depending on whether the substrate is GDP or GTP, catalyzes the formation of ppGpp or pppGpp *via* a ribosome associated mechanism (11, 12). SpoT-mediated stringent responses occur under other nutritional stresses such as fatty acid starvation (13), carbon source starvation (14), phosphorous limitation (15, 16) and iron limitation (17). In the case of carbon source diauxic growth transitions, there is evidence that RelA and SpoT-mediated responses are both involved (18). (p)ppGpp, the alarmone these enzymes synthesize, acts globally– directly and indirectly–on replication, transcription, translation (19) and protein activities (20, 21). In addition to RelA and SpoT, the pppGpp pyrophosphatase GppA also modulates intracellular levels of (p)ppGpp by converting pppGpp into ppGpp. To date, the physiological role of pppGpp remains unclear as it has been shown to be a less potent regulator than ppGpp (20, 22, 23).

It is well established that basal levels of (p)ppGpp control growth by modulating the number of ribosomes (3, 24). The sudden accumulation of (p)ppGpp provokes a quasi-immediate inhibition of growth (25) and protein synthesis (26, 27). The nature (transient, reversible) and the potency of (p)ppGpp interactions with ribosome-associated GTPases may explain how (p)ppGpp build-up contributes to slowing down growth and reduces translational activity (20). Remarkably, the ability of bacterial cells faced with nutritional insult to resume growth and recover the pre-disturbance rate, and the role of (p)ppGpp in promoting this resilience, have not been systematically studied.

In this study therefore, we investigated i) the ability of *E. coli* to cope with severe growth inhibition and ii) the contribution of (p)ppGpp metabolism to *E. coli*’s adaptive capacity to disruptions such as these. We triggered stringent response using serine hydroxamate (SHX), a serine analogue known to have this effect in *E. coli* (10). SHX addition promotes (p)ppGpp accumulation and provokes growth arrest (28, 29), because of presumed competitive inhibition with serine binding to seryl-tRNA synthetase (28, 30). The response to SHX-induced homeostasis disruption was dissected by analyzing growth rate dynamics and quantifying the intracellular levels of ppGpp and pppGpp in perturbation experiments using the WT K-12 strain of *E. coli* and Δ*relA* and Δ*gppA* mutants. Our results provide clear evidence of the resilience of *E. coli* to growth disruption caused by SHX addition and demonstrate the key role played by ppGpp–and not by pppGpp–in *E. coli*’s ability to recover its full growth capacity.

## RESULTS SECTION

### Dynamic response to SHX addition

The macroscopic effects of SHX addition were characterized in *Escherichia coli* K-12 MG1655 cells growing exponentially in M9 medium (31) with 110 mM glucose (20 g·l^−1^) as sole carbon source. Growth was performed aerobically under controlled conditions in bioreactors. SHX (0.8 mM) was added when the OD reached 3.5.

As expected, adding SHX immediately led to severe growth inhibition (Fig. 1A), with the concomitant decreases in the OUR and CER (Fig. 1B) indicating a strong decrease in respiratory activity. Combined with the interruption of acetate production (Fig. 1C) and, to a lesser extent, in glucose consumption (Fig. 1D), these events reflect a sharp reduction in metabolic activity. However, the inhibition of growth and reduction in metabolic activity were transient and about 1.5 h after SHX addition, the biomass concentration, respiratory activity and acetate production began to increase once again (Fig. 1A).

**FIG 1:**
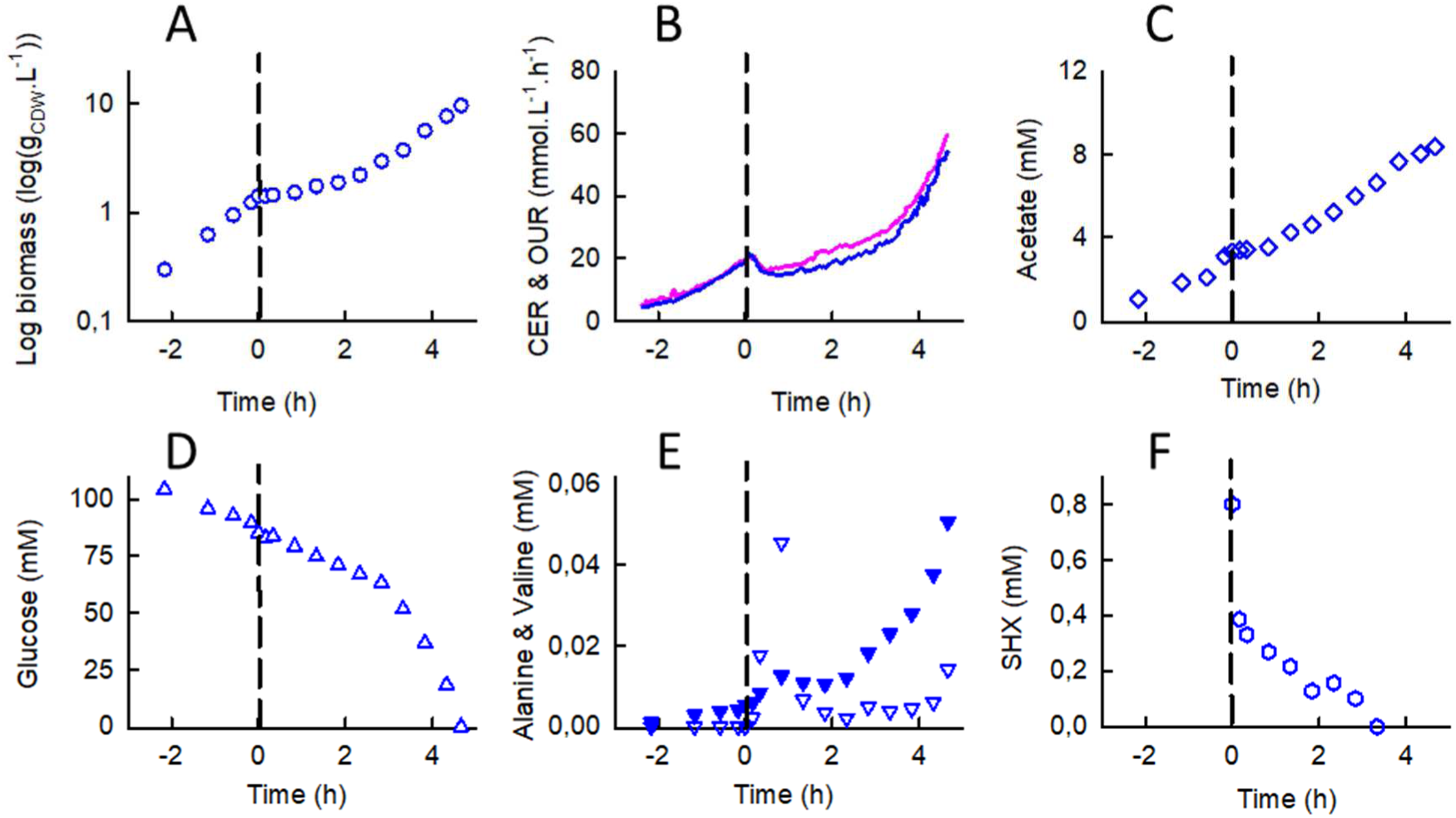
Dynamics of the response of *E. coli* K-12 MG1655 to SHX addition during exponential growth in a bioreactor: time-evolutions of (A) biomass, (B) the oxygen uptake rate (OUR, pink) and carbon dioxide evolution rate (CER, blue), (C) the acetate concentration, (D) the glucose concentration, (E) alanine (downward triangles) and valine (filled downward triangles) concentrations, and (F) the SHX concentration. Time zero was defined as the moment SHX (0.8 mM) was added to the bioreactor. The OUR and CER were determined from online measurements of O_2_, CO_2_ and N_2_ percentages as described in the Materials and Methods section. Biomass and extracellular metabolite concentrations were measured in culture samples extracted every 10 min to 1 h. The data are highly reproducible (see Fig. S1).

During the exponential growth phase, we detected relatively low levels (µM range) of alanine and valine a in the culture medium (Fig. 1E) along with other metabolic by-products (orotate, dihydroorotate – data not shown) and traces of leucine (data not shown). Interestingly, SHX addition led to a sharp increase in the concentrations of alanine and valine, which were respectively about 8 times and 2 times higher 20 min after SHX addition than just before and peaked 50 min after SHX addition (at 20 times and 3 times their pre-SHX levels, respectively). In this period, the estimated fluxes of alanine and valine excretion were respectively 9% and 2% of the fluxes required to support growth during the exponential phase (Table S1). The alanine and valine concentrations in the medium then dropped and finally increased once again as growth resumed as it was before SHX addition.

Surprisingly, we observed that the SHX concentration in the cultivation medium decreased immediately after it was added, becoming undetectable after 2.8 h (Fig. 1F). The disappearance of SHX is biotic in origin since the SHX concentration did not decrease under similar conditions but without cells (Fig. S2). In addition, NMR did not detect any of SHX degradation. To our knowledge, this has never previously been reported in literature, despite SHX being widely used to trigger the stringent response.

### *Escherichia coli* is resilient to SHX addition

Although the addition of SHX profoundly perturbs its metabolism, these results reveal that *E. coli* recovers its growth capacity, at least partially. To better characterize *E. coli’s* resilience to SHX addition, we calculated the instantaneous growth rate throughout the experiment (Fig. 2A). The growth rate before SHX addition was 0.71 ± 0.01 h^−1^ (Fig. S4A), in agreement with previously published data on K-12 *E. coli* growing in minimal medium (32, 33). The growth rate decreased suddenly down to near zero after SHX was added, before increasing constantly and levelling off about 4 h after SHX addition at a value similar to the one measured before SHX addition. Interestingly, the growth profiles measured here are typical of resilient-engineered systems able to recover their initial performance levels after disruptive events (34, 35). Here, growth is the biological property *E. coli* is able to recover. In keeping with the concept of resilience engineering, we defined three metrics to describe the resilience of *E. coli* to the disruptions caused by SHX addition: i) robustness, defined here as the residual growth rate after SHX addition, ii) the recovery rate, which is the recovery profile of the growth rate (assumed here to be linear), and iii) the recovered steady-state, which corresponds here to the growth rate at the end of the recovery process (expressed relatively to the initial growth rate). These metrics are represented by dashed lines in Fig. 2A and the corresponding numerical values are given in Table 1.

**Table 1:** Metrics quantifying the resilience of *E. coli* to SHX-induced growth disruption.

|  | Robustness (%)<sup>a</sup> | Recovery rate (h<sup>-2</sup>) | Recovered steady-state (%)<sup>a</sup> |
| --- | --- | --- | --- |
| WT | 2.8 ± 2 | 0.1944 ± 0.0201 | 96.7 ± 3.3 |
| $\Delta relA$ | 5.4 ± 2* | 0.0847 ± 0.0261 | ND<sup>b</sup> |
| $\Delta gppA$ | 10.5 ± 0.5** | 0.1941 ± 0.0482 | 98.0 ± 12.2 |

Results are presented as mean ± standard deviation (n = 3). CDW, cell dry weight.
<sup>a</sup> The robustness and recovered steady-state are expressed relative to the initial growth rate
<sup>b</sup>Not determined (only one of the three biological replicates had recovered its initial growth rate 5.5 h after SHX addition).
\* P value = 0.188
\*\* P value = 0.01332

**FIG 2:**
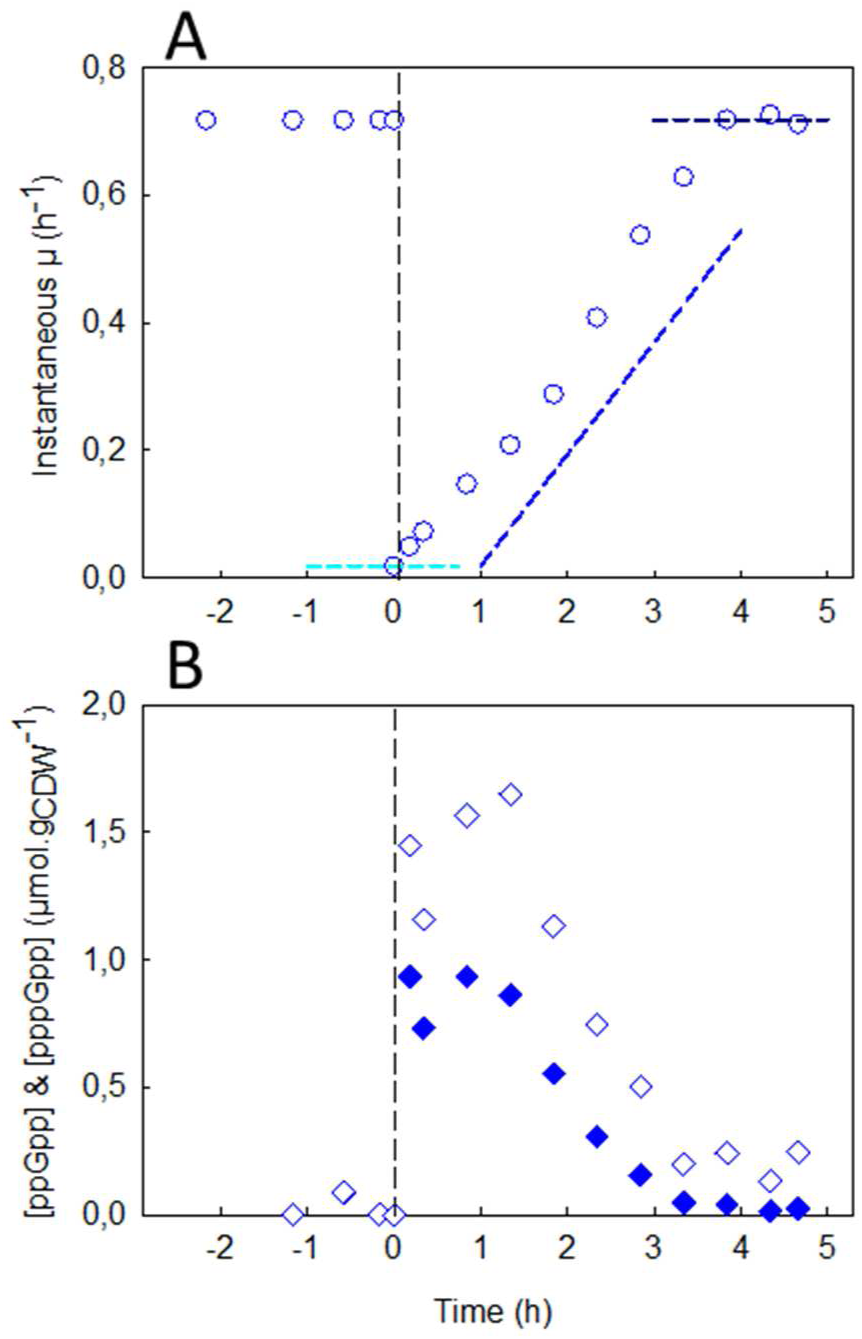
Resilience of *E. coli* K-12 MG1655 to growth disruption. (A) Instantaneous growth rate µ(t) (blue circles) as a function of time before and after SHX addition; the cyan dashed line indicates the robustness parameter, the slope of the blue dashed line is the recovery rate and the dark-blue dashed line represents the recovered steady-state. The values of these parameters are given in Table 1. (B) Intracellular concentration of ppGpp (diamonds) and pppGpp (filled diamonds) (µmol·g_CDW_^−1^) before and after addition of SHX. The data are highly reproducible (Fig. S3).

### Dynamics of intracellular (p)ppGpp levels

The intracellular concentrations of ppGpp and pppGpp were measured throughout the experiment by LC-MS/MS (36) (Fig. 2B). As expected, the addition of SHX led to a sudden intracellular accumulation of ppGpp, whose concentration was measured to be 1.45 µmol·g _CDW_ ^−1^ just a few minutes after SHX addition. The maximal concentration (1.65 µmol·g _CDW_^−1^) was reached approximatively 1 h after SHX addition. The ppGpp concentration then decreased continuously, tending toward the basal value measured during the exponential phase. Interestingly, the pppGpp concentration followed a similar profile, albeit at lower levels; the intracellular concentration of pppGpp peaked at 0.94 µmol/g_CDW_ within 1 h of SHX addition.

Because the concentrations of extracellular SHX and of intracellular (p)ppGpp both decrease during the recovery phase, it is difficult to distinguish between their respective contributions to growth recovery. Therefore, to verify that this phenomenon was associated with the stringent response and was not the result of SHX disappearing from the medium, the experiment was repeated with a Δ*relA* mutant. In *Bacillus subtilis*, deleting the *relA* gene has been reported to suppress the accumulation of (p)ppGpp in response to SHX (37).

### The stringent response is crucial for growth recovery

Before SHX addition, the growth rate of the Δ*relA* mutant was slightly lower than that of the wild-type strain (0.63 ± 0.05 h^−1^, Fig. 3A, Fig. S4A). The addition of SHX also interrupted growth and led to a reduction in metabolic activity (Supplementary Data). Note that for the Δ*relA* mutant, we did not measure pppGpp concentrations during the experiment, only those of ppGpp (Fig. 3B). The concentration of ppGpp in the exponential phase did not exceed 20 ± 3 nmol·g_CDW_^−1^, lower than the value measured for the WT strain (123 ± 85 nmol·g_CDW_^−1^) (Fig. S4B) and close to the detection limit. The presence of ppGpp in this Δ*relA* mutant indicates that under these conditions, SpoT synthesizes low levels of ppGpp in the exponential regime. As expected, we did not detect any transient accumulation of ppGpp after SHX addition. This confirms that the synthetase activity of SpoT is mainly silent in this situation and that, in agreement with previous studies (22, 26), no other RSH is involved in this response in *E. coli*. More importantly, this means that SHX addition induces growth inhibition by itself, without (p)ppGpp. Finally, although SHX disappeared completely from the medium in less than 3 h, as also observed for the WT strain (Fig. 3C), the cells had failed to fully recover their initial growth rate 6 h after SHX addition (Fig. 3A). The recovery rate of the Δ*relA* mutant was a factor of 2 lower (0.0847 ± 0.0261 h^−2^) than the WT’s (Table 1). These results highlight the crucial role of the stringent response in *E. coli*’s ability to overcome the growth disruption caused by SHX. Based on the analysis of instantaneous growth rates, WT and the Δ*relA* strains had similar robustness. Furthermore, alanine and valine were also found to accumulate in the culture medium with the Δ*relA* mutant, and the accumulation of alanine was even more pronounced with the mutant than it was with the WT strain (Table S1), indicating that this phenomenon is not related to the stringent response.

**FIG 3:**
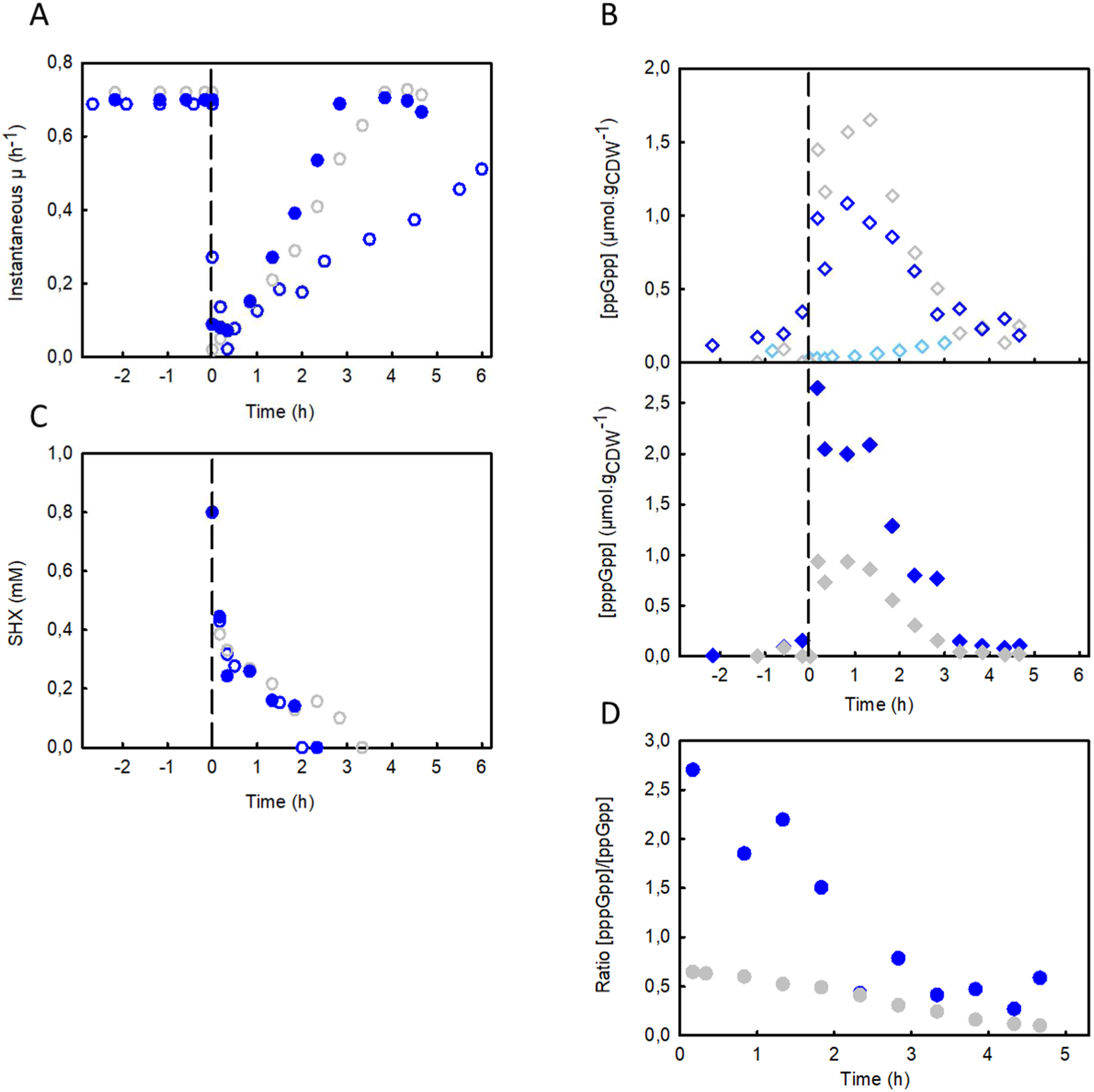
Importance of ppGpp for *E. coli*’s resilience. (A) Instantaneous growth rate µ(t) as a function of time before and after SHX addition of the Δ*relA* mutant (blue circles), the Δ*gppA* mutant (filled blue circles) and the WT (grey circles). (B) Intracellular concentrations of ppGpp (diamonds) and pppGpp (filled diamonds) (µmol·g_CDW_^−1^) before and after addition of SHX, for the Δ*relA* mutant (light blue diamonds), the Δ*gppA* mutant (blue diamonds) and the WT (grey diamonds). (C) Time evolution of the extracellular concentration of SHX (mM) for the Δ*relA* mutant (blue circles), the Δ*gppA* mutant (filled blue circles) and the WT strain (grey circles). (D) Time evolution of the [pppGpp]/[ppGpp] ratio after SHX addition for the Δ*gppA* mutant (filled blue circles) and the WT strain (filled grey circles). The corresponding results for the biological replicates are shown in Figs. S5 and S6.

### pppGpp over-accumulation has no effect on growth recovery

The physiological role of pppGpp in the stringent response in *E. coli* has remained rather unclear to date. To explore its effect on growth recovery, we applied our methodology to a mutant deleted for the *gppA* gene, which encodes for the enzyme, pppGpp 5’-gamma phosphohydrolase, which converts pppGpp to ppGpp. The Δ*gppA* mutant is known to accumulate high concentrations of pppGpp after SHX addition (23).

As expected therefore, the intracellular levels of pppGpp in the Δ*gppA* mutant were higher during the exponential phase than in the WT strain while intracellular ppGpp levels were similar (Fig. S4B). In the Δ*gppA* mutant, the concentrations of pppGpp and ppGpp were thus similar, as reported previously (23), with an estimated ppGpp/ppGpp ratio of 0.62 ± 0.38. Although the pppGpp concentration was about one order of magnitude higher than in the WT strain, the growth rates were almost identical (Fig. 3A), suggesting that pppGpp does not affect the growth rate.

As in the WT strain, adding SHX triggered the accumulation of ppGpp and pppGpp. However, while the ppGpp concentration varied around the same levels as measured for the WT strain, the pppGpp concentration was roughly twice as high as in the WT (Fig. 3B, D). The total concentration of ppGpp and pppGpp was therefore significantly higher, with the pentaphosphate form predominating, contrary to what is observed in other conditions. In this strain, the principal product of RelA is therefore pppGpp. The pppGpp/ppGpp ratio decreased over time (Fig. 3D), tending toward the value measured before SHX addition. Although this ratio is markedly different in the Δ*gppA* mutant, the growth recovery profile of the mutant was similar to that of the WT strain (with an estimated recovery rate of 0.1941 ± 0.0482 h^−2^ for the Δ*gppA* mutant; Fig. 3A and Table 1). This means that the build-up of pppGpp has no significant effect on growth recovery, the only difference in this strain being a slightly greater robustness (Table 1). Note that alanine and valine accumulated in the culture medium once again, as observed for the WT strain (Table S1).

### The decrease in (p)ppGpp concentration can be explained by growth

As shown above for the WT and Δ*gppA* strains, the addition of SHX leads to a rapid accumulation of ppGpp and pppGpp (in a few minutes), a plateau stage that last for less than one hour, and then a slow decrease in the concentrations of (p)ppGpp. It takes about 3 h in the latter phase for the intracellular concentrations of ppGpp and pppGpp to drop to the levels measured in the exponential phase (Fig. 2B). The question then arises whether the decrease in the concentration of (p)ppGpp is the result of an active degradation process or simply due to growth-driven dilution as suggested by the ppGpp and pppGpp levels being correlated with growth during this phase. To answer this question, we first calculated what the intracellular concentrations of ppGpp and pppGpp would be if they were only diluted by growth. This would require that the formation fluxes of ppGpp (*via* RelA or GppA for instance) be equal to its degradation fluxes (*via* SpoT for instance) and likewise for pppGpp, its formation flux *via* RelA being equal to its degradation fluxes *via* GppA and SpoT. In the case of the Δ*gppA* mutant, the absence of GppA makes the situation easier to evaluate. In Figs 4A and 4B, the solid black lines are the intracellular levels of ppGpp or pppGpp during the recovery phase considering growth dilution only. Because this line fits the measured intracellular concentrations of ppGpp and pppGpp relatively well, this means that the formation and degradation fluxes are equal and shows that in this strain, the decrease in ppGpp and pppGpp levels is mainly due to growth dilution rather than an active degradation process. In contrast, the measured intracellular levels of ppGpp and pppGpp do not follow this line for the WT strain (Figs 4C and 4D), meaning that a degradation process is involved. To estimate its contribution, we calculated what the intracellular ppGpp and pppGpp concentrations would be if the degradation flux were 1 to 8 times the growth dilution rate (grey lines in Fig. 4). For pppGpp (Fig. 4D), most of the experimental points are located between the first and the second grey lines, indicating that the degradation flux–likely via GppA–is about twice the growth dilution rate. The flux of ppGpp degradation is even more modest since the experimental points fall mostly between the black line and the first grey line (Fig. 4C). Altogether, these results indicate that the decrease in the concentration of (p)ppGpp is mostly accounted for by growth, even if GppA appears to participate somewhat in the degradation of pppGpp. The surprising implication of these results is that SpoT plays only a minor role in hydrolyzing ppGpp and pppGpp under these conditions.

**FIG 4:**
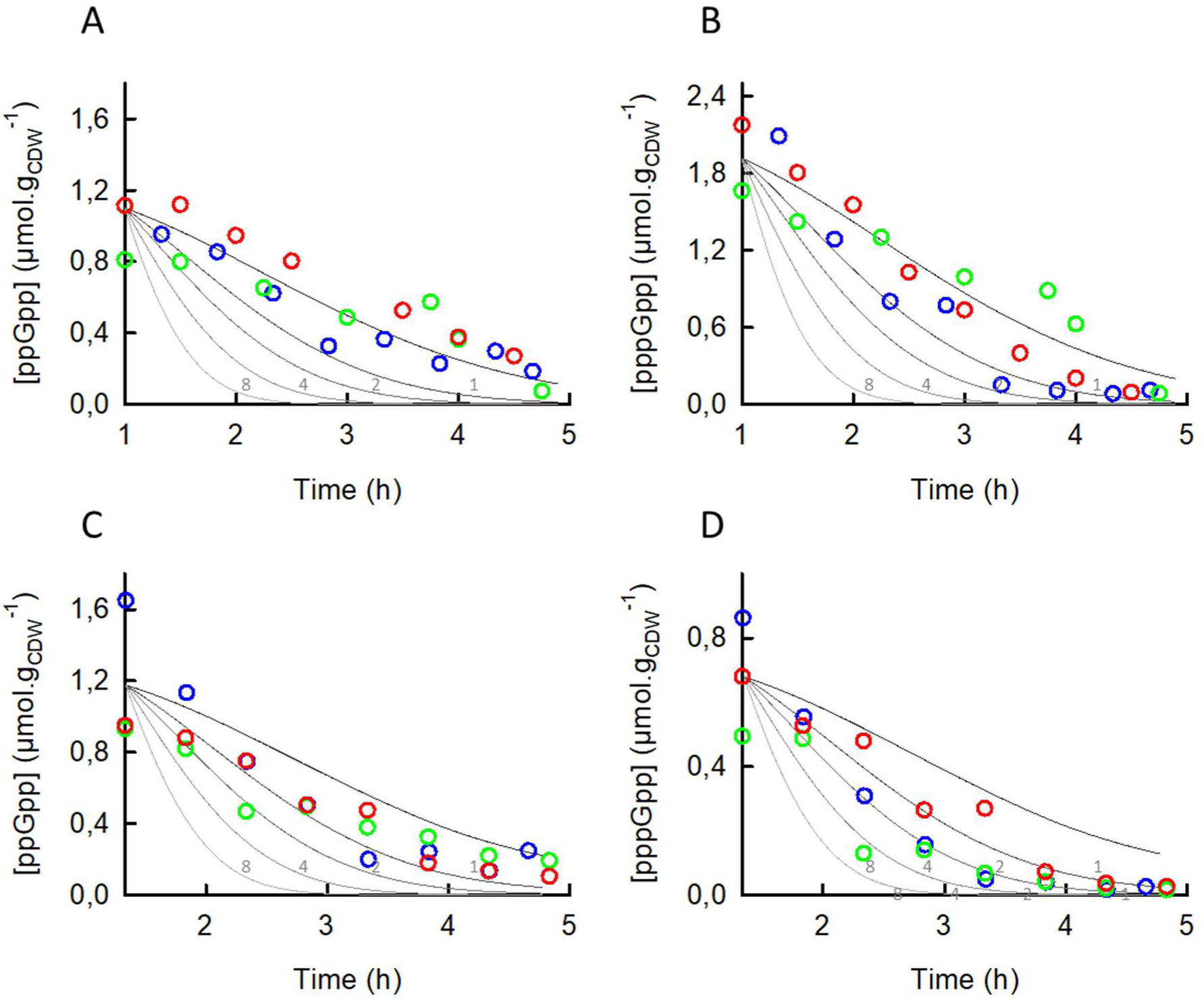
Dilution by growth of the intracellular concentrations of ppGpp and pppGpp (A, B) for the Δ*gppA* mutant and (C, D) for the WT strain of *E. coli*. In all panels, the solid lines are the concentrations of (p)ppGpp calculated using the following equation: d*(p)ppGpp*/d*t*=J_F_-J_D_-µ(t).*(p)ppGpp* where *(p)ppGpp* is the amount of ppGpp or pppGpp; J_F_ and J_D_ are respectively the fluxes of (p)ppGpp formation and degradation; and µ(t) is the instantaneous growth rate. The black lines were calculated with J_F_=J_D_, the grey lines with (J_F_-J_D_)=*a*.(µ(t).*(p)ppGpp*) with *a* ranging from 1 to 8, using as initial conditions the mean values of the (p)ppGpp concentrations and µ(t) calculated from the three biological replicates of each strain. µ(t) was calculated from the recovery rates determined for each strain, as listed in Table 1. Zero time on these graphs corresponds to the moment at which the ppGpp and pppGpp concentrations started to decline *i*.*e*. 1 h after SHX addition for the Δ*gppA* mutant and 1 h 20 min after SHX addition for the WT. The mean values of the concentrations of ppGpp and pppGpp measured in three independent repeats (blue, green and red) are plotted on panels A and B for the Δ*gppA* mutant and on panels C and D for the WT strain.

## DISCUSSION

In this study, we investigated the dynamic response of *E. coli* to severe growth disruption and the role of the stringent response in this bacterium’s ability to recover growth. To this aim, we monitored the growth and quantified consumed and excreted metabolites and intracellular levels of ppGpp and pppGpp in *E. coli* cultures before and after adding SHX.

The results demonstrate first that the K-12 WT strain of *E. coli* is resilient to SHX-induced growth disruption since its growth rate returned to pre-SHX levels a few hours after the perturbation. This recovery was first shown in the pioneering work of Tosa and Pizer (28), where growth inhibition was released by adding serine. In our work, *E. coli* appeared to recover growth by itself. Intriguingly, we observed that SHX disappeared rapidly from the medium and identified the cause as being a cell-related process, but it remains unclear whether SHX was degraded or simply internalized into the cells. The results of the experiment with the Δ*relA* mutant of K-12 *E. coli* indicate that resumption of growth is conditioned on the stringent response. Although just as with the WT, SHX disappeared from the medium, the Δ*relA* mutant failed to fully recover its pre-SHX growth rate, indicating that the stringent response is major determinant of *E. coli* ‘s resilience to growth disruption. We also observed that this resilience is not affected by an over-accumulation of pppGpp. By eliminating GppA, we inverted the pppGpp/ppGpp concentration ratio but the Δ*gppA* mutant’s recovery from SHX addition was nevertheless similar to that of the WT. Although the robustness of the Δ*gppA* mutant was slightly higher, these results indicate that pppGpp does not play a significant role in growth recovery. This is in keeping Mechold et al.’s conclusion that pppGpp is a less potent growth regulator than ppGpp (22).

The concentrations of pppGpp and ppGpp both peaked rapidly after the addition of SHX (in less than 10 min), which is in line with an earlier study of a different bacterium (38). As mentioned above, in the WT strain, the concentrations of ppGpp were higher than those of pppGpp. This is in agreement with a previous report (23) and can be explained by the activity of RelA. In the absence of *gppA* indeed, pppGpp was the dominant form in the first 2.5 h after SHX addition, as has also been reported by Mechold et al. (22). In line with these authors’ interpretation (22), this suggests that RelA may favor the synthesis of pppGpp over ppGpp, while GppA adjusts the level of ppGpp. This also supports the argument that the principal pyrophosphate acceptor is GTP (10), which remains a matter of debate in the literature (39).

It is well accepted that quantitative differences in intracellular concentrations of ppGpp may lead to different patterns of gene expression, low levels of ppGpp activating the Lrp regulon while high levels activate RpoS and the general stress response (26, 40). In addition, the hydrolase activity of SpoT is thought to be inhibited under physiological stress and in particular, in the presence of high levels of uncharged tRNA (14). We can therefore assume that the accumulation of ppGpp following the addition of SHX activates stress survival genes in an RpoS-dependent manner thereby inhibiting the hydrolase activity of SpoT. This inhibition would explain the plateau in the ppGpp concentration and the decrease in the (p)ppGpp levels by growth dilution rather than active degradation.

Our results also show that it is SHX itself that provokes growth arrest and that this growth arrest is associated with the excretion of alanine, valine and to a lesser extent leucine. These three amino acids are derived from pyruvate and along with glycine, are the most abundant amino acids in terms of biomass (41). Because this excretion is substantial and occurs after SHX addition, it can be interpreted as a (transient) metabolic overflow in response to a sudden drop in the demand for proteinogenic amino acids. However, further investigations are required to elucidate the regulatory mechanism underlying this metabolic overflow.

### Concluding remarks

This report promotes a better understanding of the resilience of *E. coli* to severe growth disruption and the role of (p)ppGpp metabolism in this phenomenon. Our results and data, specifically the ppGpp and pppGpp concentrations, will hopefully serve as a hypothesis-generating resource for future studies on (p)ppGpp metabolism and more generally on the stringent response, a crucial process in bacterial adaptation and survival.

## EXPERIMENTAL SECTION

### Chemicals and reagents

DL-serine hydroxamate (SHX) was purchased from Sigma Aldrich (St. Quentin-Fallavier, France). LC-MS grade solvents (methanol, acetonitrile) were obtained from Instrumentation Consommables et Service (ICS, Lapeyrousse-Fossat, France).

### Bacterial strains and growth conditions

All strains were derived from *E. coli* strain K-12 MG1655. The Δ*relA* and Δ*gppA* strains were constructed by P1 transduction of gene deletions marked with a kanamycin resistance cassette from the Keio collection (42). The kanamycin resistance cassette was removed using FLP recombinase from pCP20 plasmid (43). All strains, plasmids and primers are listed in Table S2 and the genetic modifications were checked by PCR.

Cells were cultured on M9-based synthetic minimal medium supplemented with glucose (31). Glucose, thiamine and MgSO_4_ were sterilized by filtration (Minisart 0.2 μm syringe filter, Sartorius, Göttingen, Germany) and other solutions were autoclaved separately. Thiamine was added to a final concentration of 0.1 g/L. All stock cultures were stored at −80°C in lysogeny broth (LB) medium containing glycerol (40 %, v/v). For the cultures, 5 mL of overnight cultures in LB were used as inoculum and then sub-cultured in shake-flasks containing 50 mL of minimum medium with 3 g/L glucose starting at OD600nm = 0.05 and incubated at 37 °C and 210 rpm for 15 hours in an orbital shaker (Inova 4230, New Brunswick Scientific, New Brunswick, NJ, USA). Cells were harvested during the exponential growth phase by centrifugation for 10 min at 10,000 *g* at room temperature with a Sigma 3-18K centrifuge (Sigma Aldrich, Seelze, Germany), washed with the same volume of fresh medium (without glucose or thiamine), and used to inoculate 500 mL bioreactors (Multifors, Infors HT, Bottmingen, Switzerland) containing 300 ml of minimal medium with 20 g/L glucose (110 mM) at OD600nm = 0.15. The temperature was set to 37°C and the pH was maintained at 7 by automatically adding 14% (g/g) ammonia or 11 % (g/g) phosphoric acid. Aeration and the stirrer speed were controlled to maintain adequate aeration (DOT > 30% saturation). Cell growth was monitored by measuring the optical density at 600 nm with a Genesys 6 spectrophotometer (Thermo, Carlsbad, CA, USA) The percentages of O_2_, CO_2_ and N_2_ concentrations were measured in the gas output during the culture process using a Dycor ProLine Process mass spectrometer (Ametek, Berwyn, PA, USA), and the data obtained was used to calculate the oxygen uptake rate (OUR) and carbon dioxide emission rate (CER). Stringent response was triggered by adding SHX at 0.8 mM to the culture when the OD reached 3.5.

### Calculation of the instantaneous growth rate

The instantaneous growth rate (µ_(t)_) was determined by fitting the time evolution of the biomass concentration i) to an exponential function prior to SHX addition and ii) to a parametric function after SHX addition, from which µ_(t)_ was calculated as µ_(t)_ = dX/(X·dt).

### Sampling and (p)ppGpp extraction

Culture medium (400 µL) was withdrawn from the bioreactor and vigorously mixed with 4.5 mL of a pre-cooled acetonitrile/methanol/H_2_O (4:4:2) solution at −40°C to rapidly quench metabolic activity (36). Immediately thereafter, 100 µL of ^13^C labeled metabolites were added to the latter mixture as internal standards. The tubes were then placed in a cooling bath of ethanol pre-cooled at −40°C and evaporated to dryness in a SpeedVac (SC110A SpeedVac Plus, ThermoSavant, Waltham, MA, USA) under vacuum for 4 h and then stored at −80°C until needed.

### IC-ESI-HRMS quantification of (p)ppGpp

ppGpp and pppGpp were quantified as described in ref. (36). Briefly, after resuspension of the cell extract samples in 20 mM ammonium acetate buffer at pH 9 to a final volume of 500 µL, cell debris were removed by centrifugation at 10,000 g for 10 min at 4°C. The samples were then analyzed using an ion chromatograph (Thermo Scientific Dionex ICS-5000+ system, Dionex, Sunnyvale, CA, USA) coupled to a LTQ Orbitrap mass spectrometer (Thermo Fisher Scientific, Waltham, MA, USA) equipped with an electrospray ionization probe. Mass spectrometry analysis was performed in the negative FTMS mode at a resolution of 30,000 (at *m*/*z* = 400) in full scan mode, with the following source parameters: capillary temperature, 350°C; source heater temperature, 300°C; sheath gas flow rate, 50 a.u. (arbitrary unit); auxiliary gas flow rate, 5 a.u;, S-Lens RF level, 60%; and ion spray voltage, 3.5 kV. The data were acquired using the Xcalibur software (Thermo Fisher Scientific, Waltham, MA, USA). Three samples from three independent biological replicates were analyzed.

### NMR analysis of culture supernatants

Exocellular metabolites were identified and quantified by nuclear magnetic resonance (NMR). Broth samples were collected at different times and filtered (Minisart 0.2 μm syringe filter, Sartorius, Göttingen, Germany). The supernatants, consisting of the culture medium, were mixed with 100 µL of D_2_O with 2.35 g/L of TSP-d4 (deuterated trimethylsilylpropanoic acid) as internal reference. Proton NMR spectra were recorded on an Avance III 800 MHz spectrometer equipped with a 5 mM QCI-P cryo probe (Bruker, Rheinstatten, Germany). Quantitative ^1^H-NMR was performed at 280 K, using a 30° pulse and a relaxation delay of 10 s. Two-dimensional ^1^H–^13^C heteronuclear single quantum coherence (HSQC) spectra were recorded at 280 K to quantify SHX in the supernatant. Sixteen scans were co-added with 4096 x 128 points and 13.35 x 60 ppm spectral widths. The spectra were processed and the metabolites were quantified using Topspin 3.1 (Bruker, Rheinstatten, Germany). Extracellular metabolites from three independent biological replicates were analyzed.

## Acknowledgments

The authors gratefully acknowledge financial support from the Bioprocess R&D department of Sanofi Pasteur and from the Association Nationale de la Technologique (ANRT). The authors also thank MetaToul (Metabolomics & Fluxomics Facilities, Toulouse, France, www.metatoul.fr) and its staff for technical support and access to NMR and mass spectrometry facilities. MetaToul is part of the French National Infrastructure for Metabolomics and Fluxomics (www.metabohub.fr), funded by the ANR (MetaboHUB-ANR-11-INBS-0010).

---

**Table S1:**
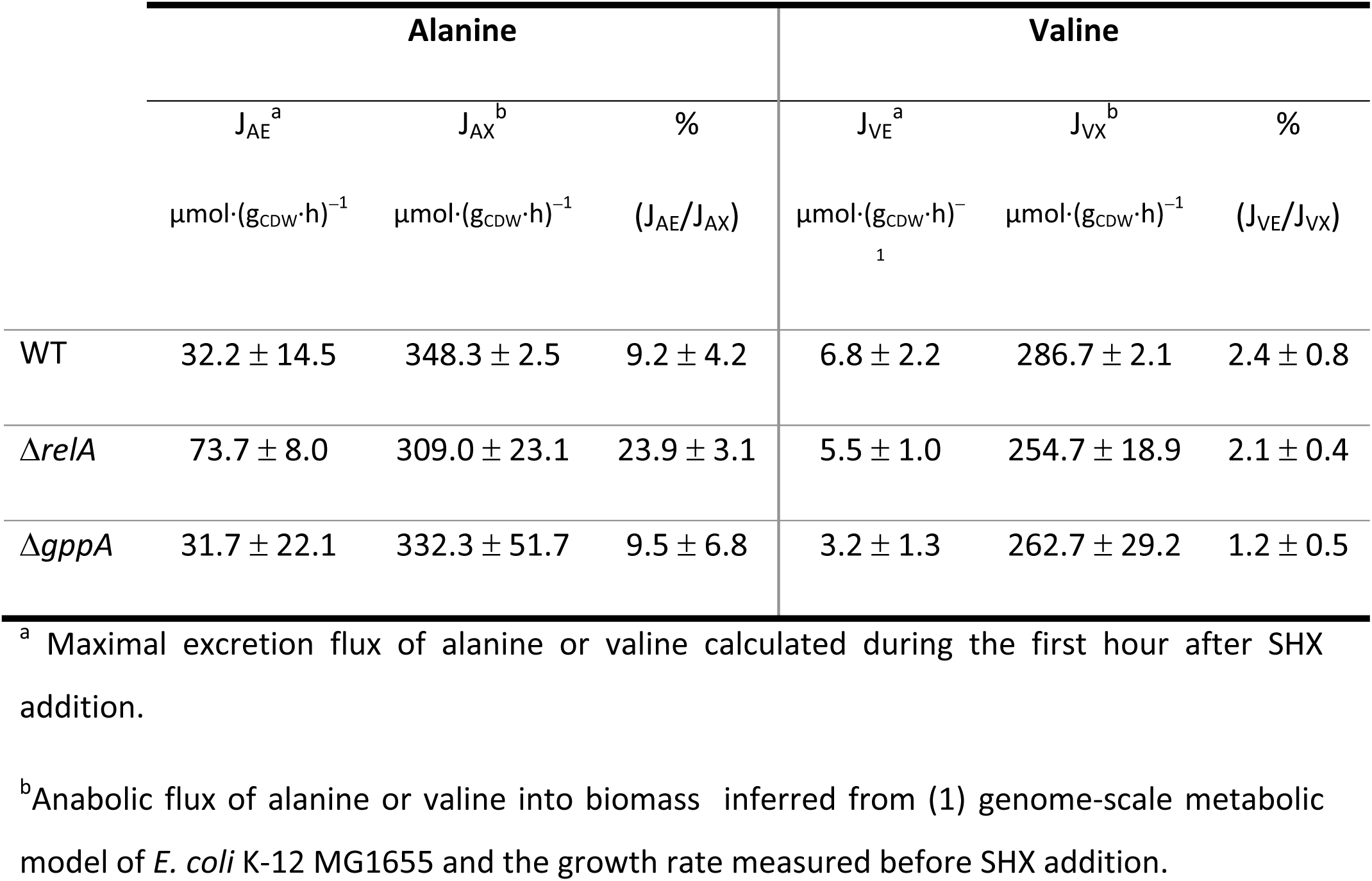
Fluxes of alanine and valine excretion following SHX addition.

|  | Alanine |  |  | Valine |  |  |
| --- | --- | --- | --- | --- | --- | --- |
| | $J_{AE}^a$ | $J_{AX}^b$ | % | $J_{VE}^a$ | $J_{VX}^b$ | % |
| | $\mu\text{mol}\cdot(\text{g}_{\text{CDW}}\cdot\text{h})^{-1}$ | $\mu\text{mol}\cdot(\text{g}_{\text{CDW}}\cdot\text{h})^{-1}$ | $(J_{AE}/J_{AX})$ | $\mu\text{mol}\cdot(\text{g}_{\text{CDW}}\cdot\text{h})^{-1}$<br>1 | $\mu\text{mol}\cdot(\text{g}_{\text{CDW}}\cdot\text{h})^{-1}$ | $(J_{VE}/J_{VX})$ |
| WT | $32.2 \pm 14.5$ | $348.3 \pm 2.5$ | $9.2 \pm 4.2$ | $6.8 \pm 2.2$ | $286.7 \pm 2.1$ | $2.4 \pm 0.8$ |
| $\Delta relA$ | $73.7 \pm 8.0$ | $309.0 \pm 23.1$ | $23.9 \pm 3.1$ | $5.5 \pm 1.0$ | $254.7 \pm 18.9$ | $2.1 \pm 0.4$ |
| $\Delta gppA$ | $31.7 \pm 22.1$ | $332.3 \pm 51.7$ | $9.5 \pm 6.8$ | $3.2 \pm 1.3$ | $262.7 \pm 29.2$ | $1.2 \pm 0.5$ |

<sup>a</sup> Maximal excretion flux of alanine or valine calculated during the first hour after SHX addition.
<sup>b</sup> Anabolic flux of alanine or valine into biomass inferred from (1) genome-scale metabolic model of *E. coli* K-12 MG1655 and the growth rate measured before SHX addition.

**Table S2:**
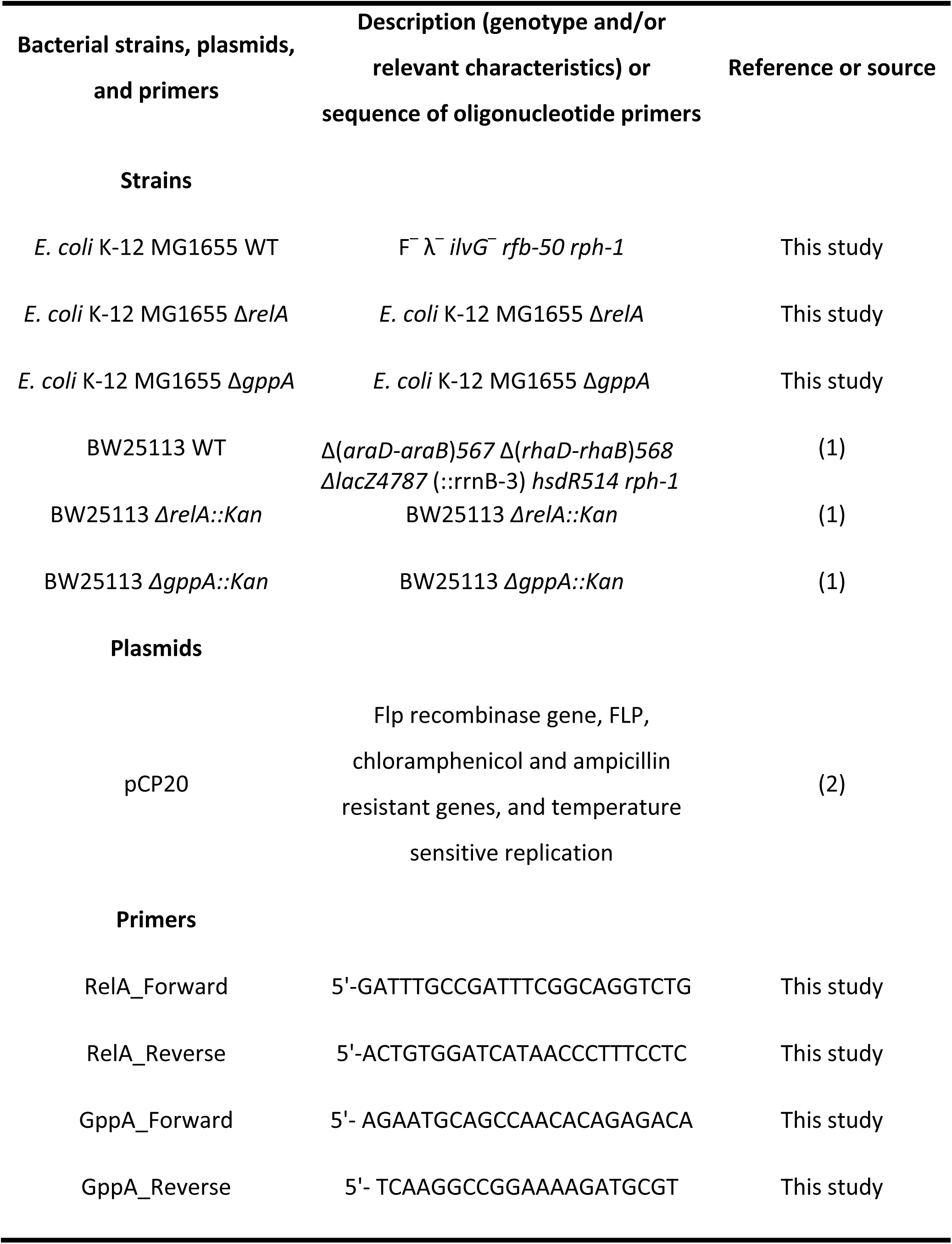
Strains, plasmids and primers used in this study.

**FIG S1:**
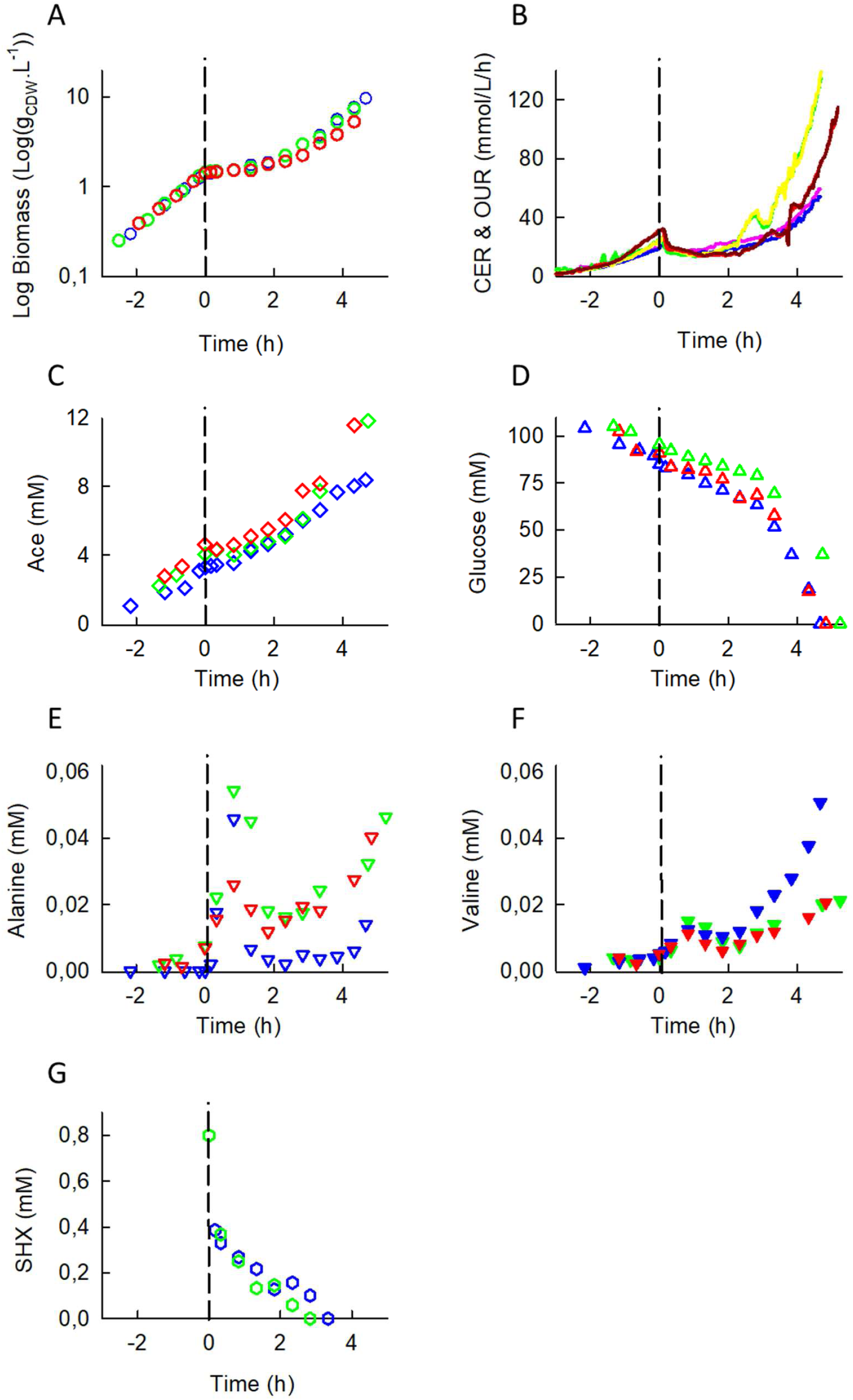
Reproducibility analysis of the response of *E. coli* K-12 MG1655 to SHX addition. (A) biomass, (B) oxygen uptake rate (OUR) and carbon dioxide evolution rate (CER), (C) acetate concentration, (D) glucose concentration, (E) alanine concentration, (F) valine concentration and (G) SHX concentration. The data from Fig. 1 are shown in blue, repeat #1, in green, and repeat #2, in red; except for the OUR (Fig. 1 data, pink; repeat #1, green; repeat #2, red) and the CER (Fig. 1 data, blue; repeat #1, yellow; repeat #2, dark red).

**FIG S2:**
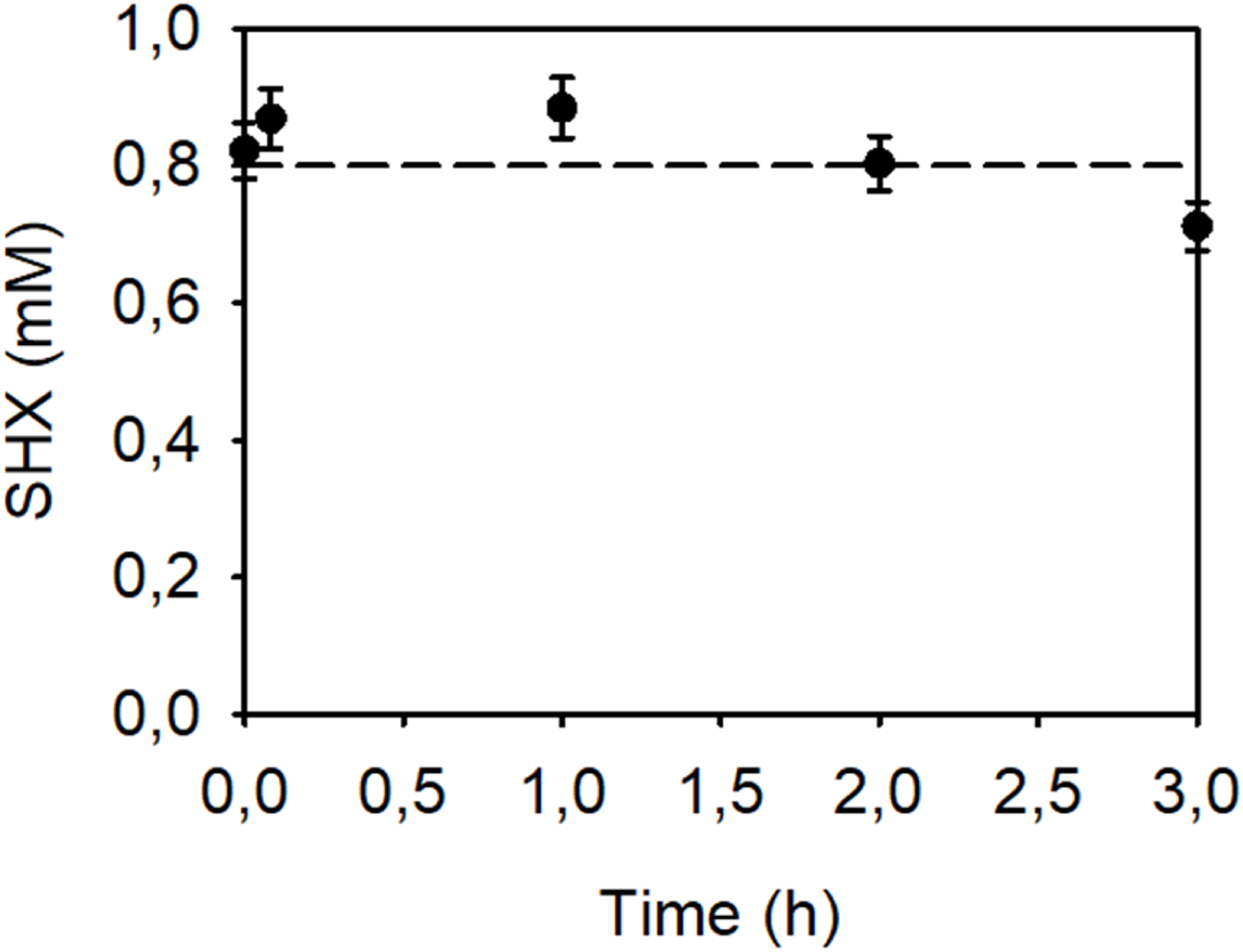
Stability of the SHX concentration over time. SHX (0.8 mM) was prepared in the culture medium (pH = 7) used in this work for the cultures of *E. coli* and incubated at 37°C. Samples (n=3) were taken and the SHX concentration was measured by NMR as described in the Materials and Methods section.

**FIG S3:**
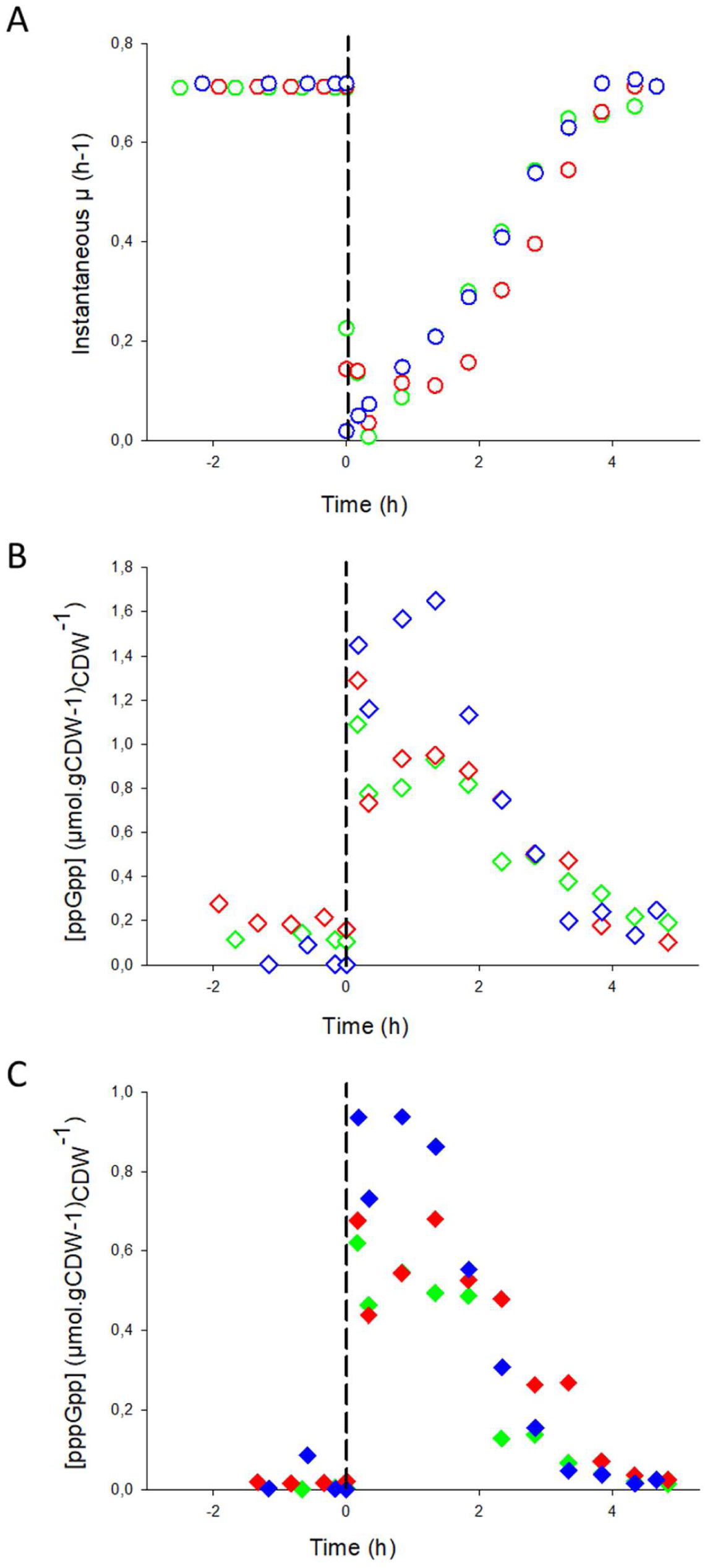
Reproducibility analysis of *E. coli* K-12 MG1655’s resilience to growth disruption. (A) Instantaneous growth rate, µ(t), and (B,C) intracellular concentrations of (B) ppGpp and (C) pppGpp. The data from Fig. 2 are shown in blue, repeat #1, in green, and repeat #2, in red.

**FIG S4:**
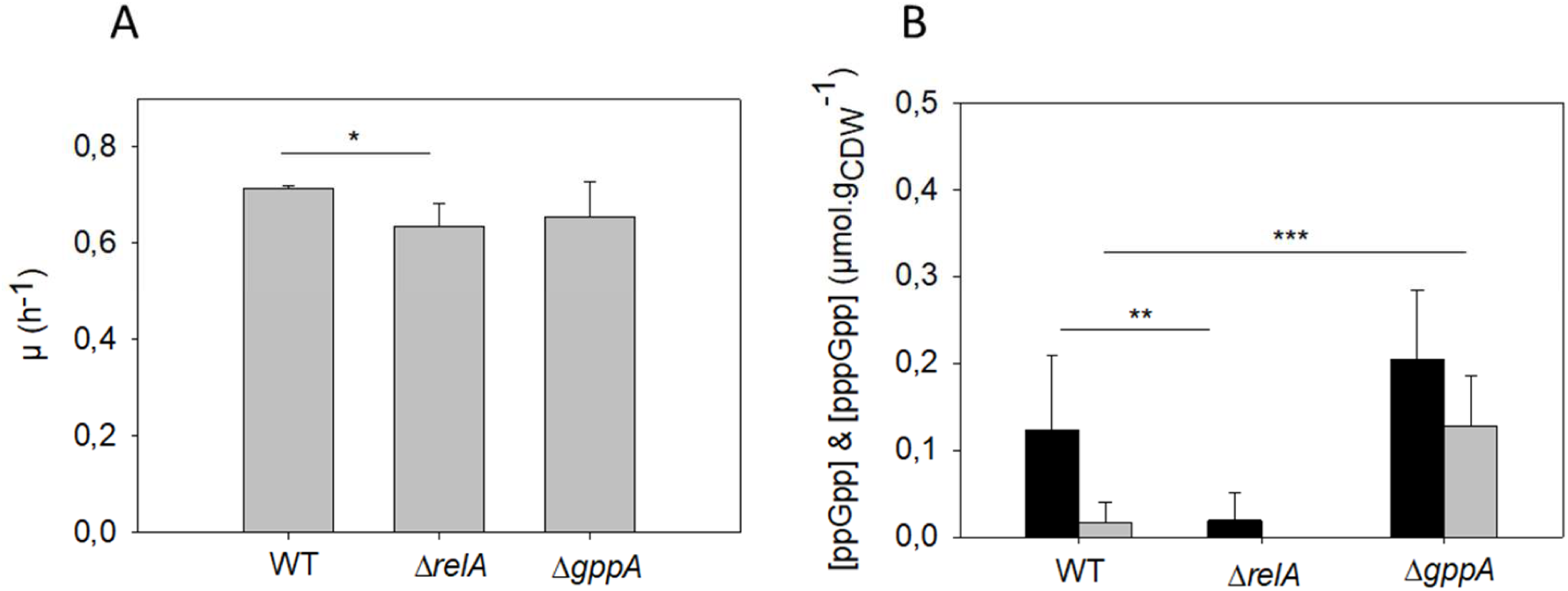
(A) Growth rate (h^−1^) in the exponential phase (before SHX addition) determined for the WT, Δ*relA* and Δ*gppA* strains of *E. coli* K-12 MG1655. The average values and standard errors of the means were calculated from the values measured in three biological replicates. (B) Intracellular concentrations of ppGpp (black bars) and pppGpp (gray bars) during the exponential phase in the WT, Δ*relA* and Δ*gppA* strains. The average values and standard errors of the means were calculated from at least 6 values measured in three biological replicates. T-test p-values: * p < 0.05, **p < 0.02, ***p < 0.005.

**FIG S5:**
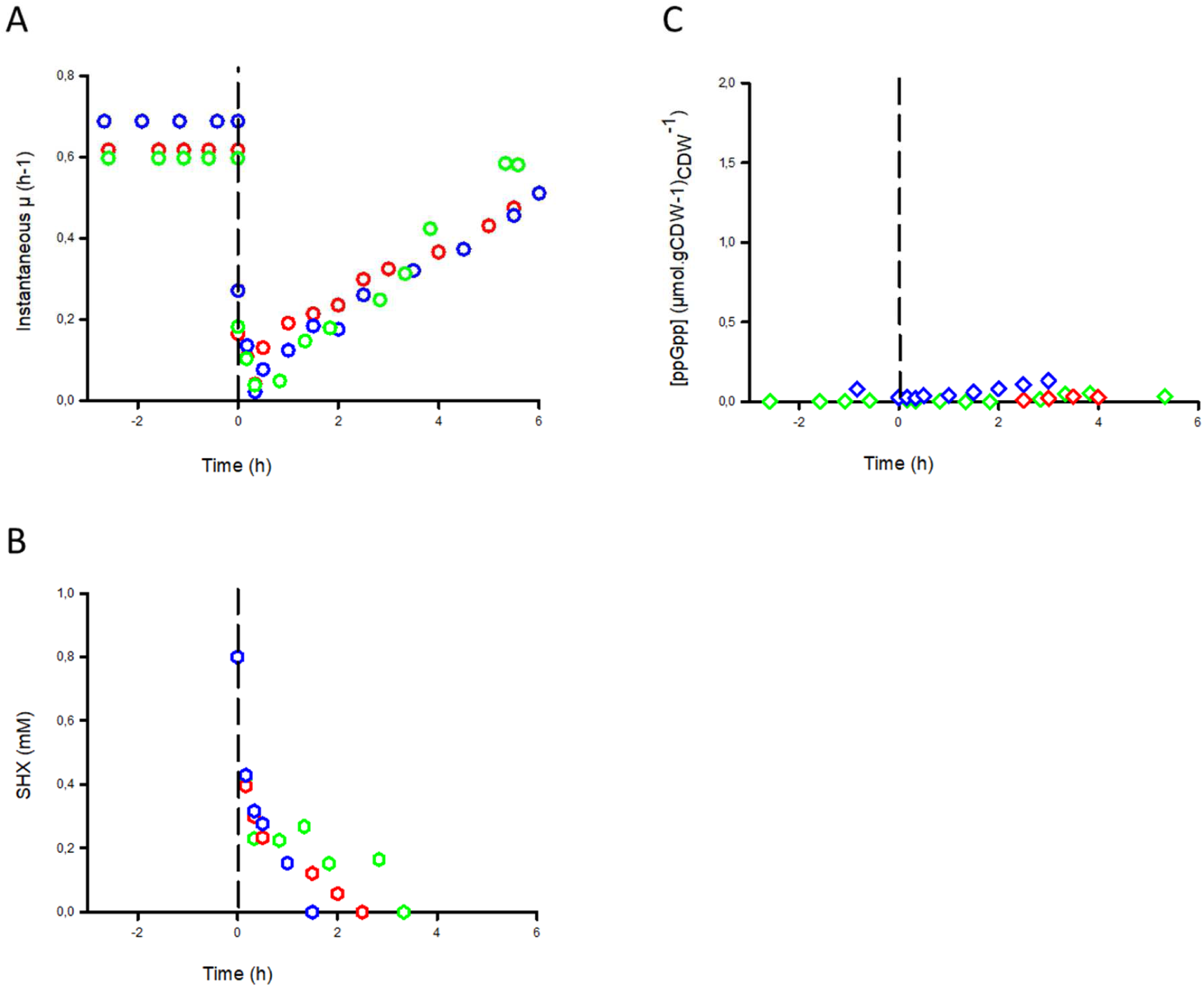
Reproducibility analysis of the response *E. coli* K-12 MG1655 Δ*relA* to SHX addition. (A) Instantaneous growth rate, µ(t), (B) SHX concentration, and (C) intracellular concentrations of ppGpp. The data from Fig. 3 are shown in blue, repeat #1, in green, and repeat #2, in red.

**FIG S6:**
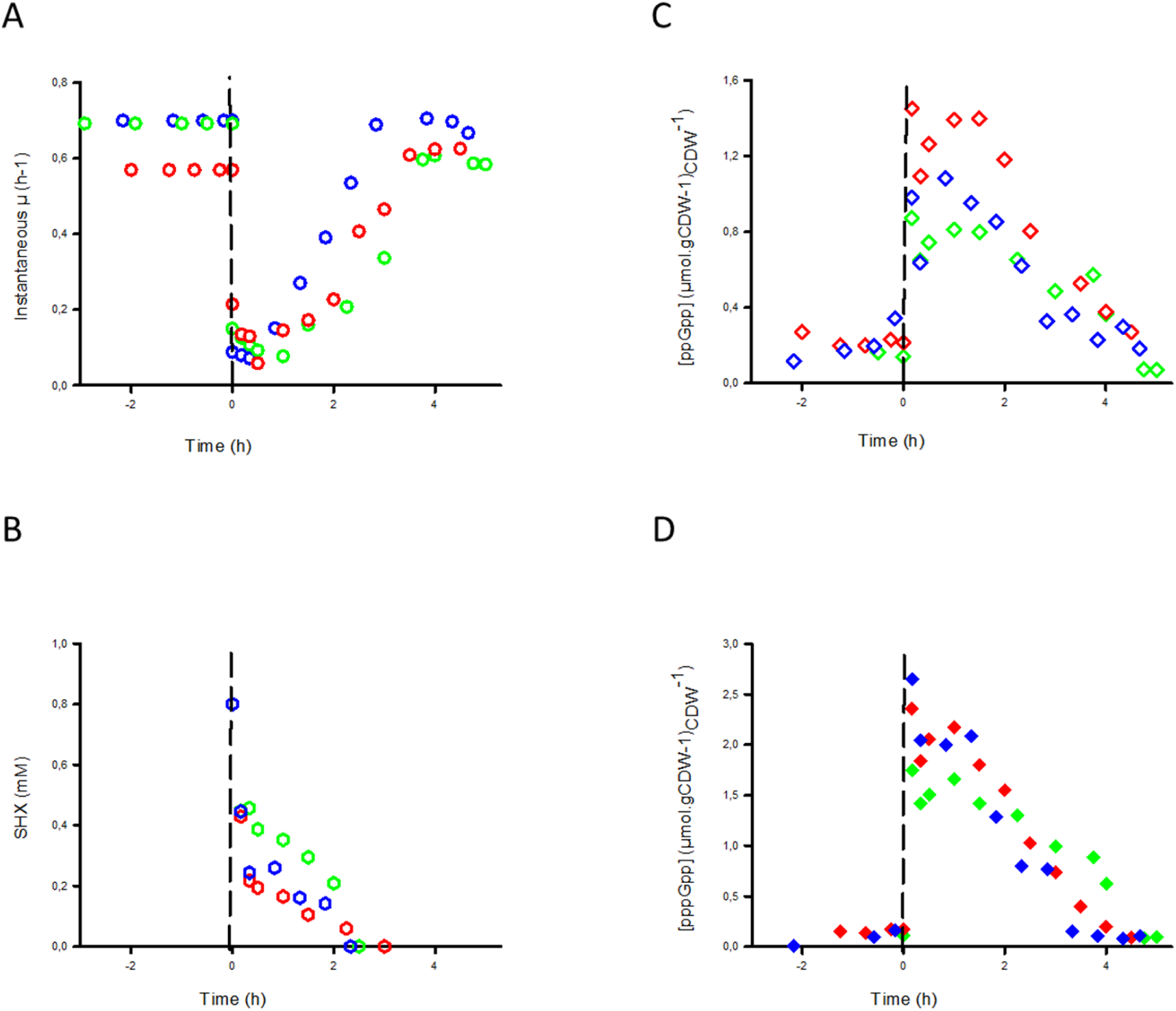
Reproducibility analysis of the response of E. coli K-12 MG1655 Δ*gppA* to SHX addition. (A) Instantaneous growth rate, µ(t), (B) SHX concentration, and (C,D) intracellular concentrations of (C) ppGpp and (D) pppGpp. The data from Fig. 3 are shown in blue, repeat #1, in green, and repeat #2, in red.

